# Upregulated GRP78 and sFlt-1 in preeclampsia induces IRE1 and ATF6 arms of UPR leading to ER stress in placental cells

**DOI:** 10.1101/2023.05.30.542817

**Authors:** Sankat Mochan, Sunil k Gupta, Pallavi Arora, Neerja Rani, Neerja Bhatla, Sada Nand Dwivedi, Renu Dhingra

**Author notes:** Correspondence: Professor (Dr.) Renu Dhingra [ ].

## Abstract

**Introduction:** Amongst vivid and diversified stresses which placenta in preeclampsia undergoes, ER stress has been a hal of the fame which is an insinuation of ill-fated UPR. In our previous study, we have already reported activation of PERK arm due to sFlt-1 present in preeclamptic mothers. The present study is an attempt to interpret rest of the two arms (IRE1 and ATF6) of UPR and ER stress subsequently by *in vitro* study using BeWo cells after upregulation of master regulator of UPR (GRP78) in placental tissue.

**Materials and Methods:** Part I: Serum analysis of circulating levels of GRP78 and sFlt-1 [in 50 pairs of preeclamptic and normotensive, non-proteinuric (control) pregnant women)] using ELISA.

Part II: Correlation analysis of levels of GRP78 and sFlt-1 in preeclamptic and control groups. Part III: Status of expression of GRP78 in placentae (n=10 each, preeclamptic and control groups) was reported using immunofluorescence.

Part IV: *In vitro* experiments using BeWo cells were carried out to analyse the effect of GRP78 and sFlt-1 on IRE1 and ATF6 arms of UPR at protein (immunofluorescence and western blot) and transcript (qRT-PCR) levels.

**Results:** Protein expressions of GRP78 and sFlt-1 were found significantly elevated in preeclamptic patients as compared to controls. Enhanced expression of master regulator of UPR (GRP78) in placental tissue of preeclamptic women was reported. Up-regulated expression of XBP1 (IRE1 arm) and ATF6 markers of UPR was observed in trophoblast cells.

**Conclusion:** The upregulated expression of GRP78 in preeclamptic placentae and enhanced expression of XBP1 and ATF6 markers in BeWo cells at both protein and transcript levels connote role played by raised circulating levels of GRP78 and sFlt-1 in preeclampsia.

## Introduction

Preeclampsia (PE), affecting 8% (nearly 10% in developing/under-developed nations) of all pregnancies worldwide and is diagnosed as emergence of hypertension (systolic blood pressure of >140 mm Hg or diastolic blood pressure of >90 mm Hg) and substantial proteinuria (≥300 mg in 24 h) at or after 20 weeks of gestation.^1,2,3,4^ Albeit its precise aetiology is unclear till date, the systematic manifestations are sought to be associated with poor placentation for which a number of pathological paradigms have been postulated such as ischemia/hypoxia, inflammation, oxidative stress, and endoplasmic reticulum (ER) stress.^5,6^ ER stress comes into role because of in-capacious unfolded protein response (UPR) to restore proteostasis and failure of glucose regulated protein78 (GRP78) to bind with trans-membrane sensors (PERK, IRE1 and ATF6) placed at luminal aspect of ER.^7,8^ Apart from the ER stress, placenta of preeclamptic women manifests upregulated sFlt-1 (soluble fms like tyrosin kinase-1, an anti angiogenic protein) at the time of disease presentation.^9,10,11,12^

So far no study is available in literature explaining ER stress in preeclamptic women in view of impact of raised central regulator of UPR along with elevated sFlt-1 levels in maternal circulation. Thus, it is in this background, our present study is an attempt to highlight the ER stress in preeclamptic placentae and cells procreated due to raised GRP78 and sFlt-1 via two independent arms of UPR, IRE1 and ATF6.

## Materials and Methods Study Subjects

100 pregnant women with singleton pregnancy visiting in-patient ward of All India Institute of Medical Sciences, New Delhi, India, were recruited for this cross sectional case control study. The preeclamptic women (n=50) after clinical diagnosis as per ACOG guidelines were enrolled as cases and normotensive, non-proteinuric pregnant women (n=50) (maternal and gestational age matched) without FGR and other medical complications were enrolled as controls. Written informed consents from all enrolled pregnant women were received after approval of study protocol from Institute Ethics Committee, AIIMS, New Delhi. Sera from the venous blood was collected, centrifuged at 1200 RPM for 4 minutes, and stored in aliquots at -80°C for ELISA and cell culture experiments. Protein levels of GRP78 and sFlt-1 were determined in serum samples of both patients and controls by ELISA. 20 caesarean delivered placentae (10 each, PE and normotensive) were used to analyze the protein expression of GRP78 by Immunofluorescence (IF) staining.

## ELISA

Sandwich ELISA (GRP 78 ELISA Kit: Enzo Life Sciences, Inc., sFlt-1 ELISA kit: R&D Systems Inc., Minneapolis, MN, U.S.A.) was done. Circulating levels of GRP78 and sFlt-1 in patients (preeclamptic) and controls (normotensive, non-proteinuric) were estimated.

### Correlation analysis

Spearman correlation coefficient was used to determine the linear correlation between GRP78 and sFlt-1.

### Immunofluorescence microscopy in placentae

Paraffin tissue blocks were sectioned on microtome (Thermo Scientific™ HM 325) and were taken on Poly-L-lysine (Sigma) coated slides. Two changes of xylene (each for 5 minutes) followed by two changes of absolute alcohol (each for 3 minutes) and subsequently one change of 90% alcohol were given for 1 minute. Slides were rinsed with distilled water. Antigen retrieval was done with sodium citrate buffer (15 min at 95-100°C) followed by treatment with TBSTx (2 min) (2 times). BSA (blocking agent) was applied on slides for 35 minutes followed by overnight incubation with primary antibody at 4°C. Slides were then rinsed with PBST20 followed by incubation with secondary antibody for 2.5 hours and then washed with PBS. Mounting was done with fluoroshield mounting media with DAPI (Abcam). Stained slides were observed under Nikon Eclipse Ti-S elements microscope using NiS-AR software. Other chemicals (analytical grade) were procured from Fischer Scientific.

### *In vitro* experiments

The human choriocarcinoma cell line (BeWo) was procured from American Type Culture Collection (ATCC) and maintained in F-12 HAM nutrient medium supplemented with 10% fetal bovine serum, 100 U/ml pencillin, 100μg/ml Streptomycin. Cells were passaged with 0.025% trypsin and 0.01% EDTA.

Depending on various treatments given to BeWo cells, study was divided into four experimental groups. Cells of group 1 were exposed to sera from normotensive mothers (NT) who had lower levels of GRP78 and sFlt-1. Group 2 cells were those, exposed to sera from preeclamptic pregnant women (PE). BeWo cells treated with tunicamycin served as positive control and were marked as group 3. The cells of group 4 did not receive any treatment. After the various treatments, induction of UPR arms [in form of activated IRE1 (XBP1) and ATF6] of ER stress pathway were studied, at various time points (8h, 14h, 24h) at protein (Immunofluorescence microscopy, Western blot) and transcript level (qRT-PCR).

### Immunofluorescence microscopy

BeWo Cells were trypsinized, seeded, allowed to grow on coverslips in multiple well chamber and incubated at 37°C in 5% CO_2_. After 8h, 14 h and 24 h, cells were taken out from incubator and washed with PBS, fixed in 4% PFA for 15 min at room temperature. After fixation, cells were washed with PBS and permeabilized with PBS + 0.1% Triton X-100 followed by PBS washing. Nonspecific blocking was done using 5% normal goat serum in PBS and Triton X. Cells were incubated for 12 hours at 4°C in primary antibodies [anti XBP1 antibody (1:200), anti ATF6 antibody (1:1000)]. Cells were washed with PBSTx and thereafter incubated in secondary antibody in 1:500 dilution for 1 hour at room temperature in dark room. Cells were washed in PBS and mounted with flouroshield mounting media with DAPI on the slide and observed under the fluorescence microscope (Nikon Eclipse Ti-S elements using NiS-AR software).

### Western blot analysis

Trophoblast Cells (BeWo) were lysed in SDS-PAGE sample buffer [10% SDS, 60 mM Tris-HCl (pH 6.8), 10% glycerol, 0.001% bromophenol blue, 0.33% mercaptoethanol] and boiled for 5 min. The lysates were analyzed by immunoblotting using 1:500 of anti XBP1 antibody and 1:1000 of anti ATF6 (Abcam) for 12 h at 4°C. The blots were then incubated in secondary antibody (HRP conjugated) for 2 hours. The blots were visualized by treating the membranes in DAB Tetrahydrochloride and H_2_O_2_. β-actin was used as protein loading control. Densitometric analysis was done subsequently and the blots were scanned thereafter in a gel documentation system, using Quantity 1 software (Bio-Rad, Hercules, CA, USA).

### qRT-PCR (quantitative Real Time-Polymerase Chain Reaction)

RNA extraction was done from treated cells via Ambion (Invitrogen) kit followed by c-DNA conversion by Thermo revert aid H-minus reverse transcriptase kit. cDNA was amplified by quantitative RT-PCR for determining mRNA expression of XBP1 and ATF6 against gene of interest with an internal control (β actin and GAPDH). Primers were designed by NCBI (Table 1).

### Statistical Analysis

Data was analyzed by STATA 14.1 and Graph Pad Prism 7. Average levels of the variables between the two groups were compared by paired t-test and Wilcoxon signed rank test. Correlation analysis was done by Spearman correlation coefficient. Relative quantification cycles of gene of interest (ΔCq) was calculated by ΔCq = Cq (target) - Cq (reference). Relative mRNA expression with respect to internal control gene was calculated by 2^-ΔCq^. For comparing more than two groups, ANOVA with Bonferroni’s multiple comparison test and Kruskal Wallis with Dunn’s multiple comparison test were used. *p* value<0.05 was considered statistically significant.

## Results

Clinical characteristics of patients and controls are summarized in Figure 1, Table 2.

**Figure 1:**
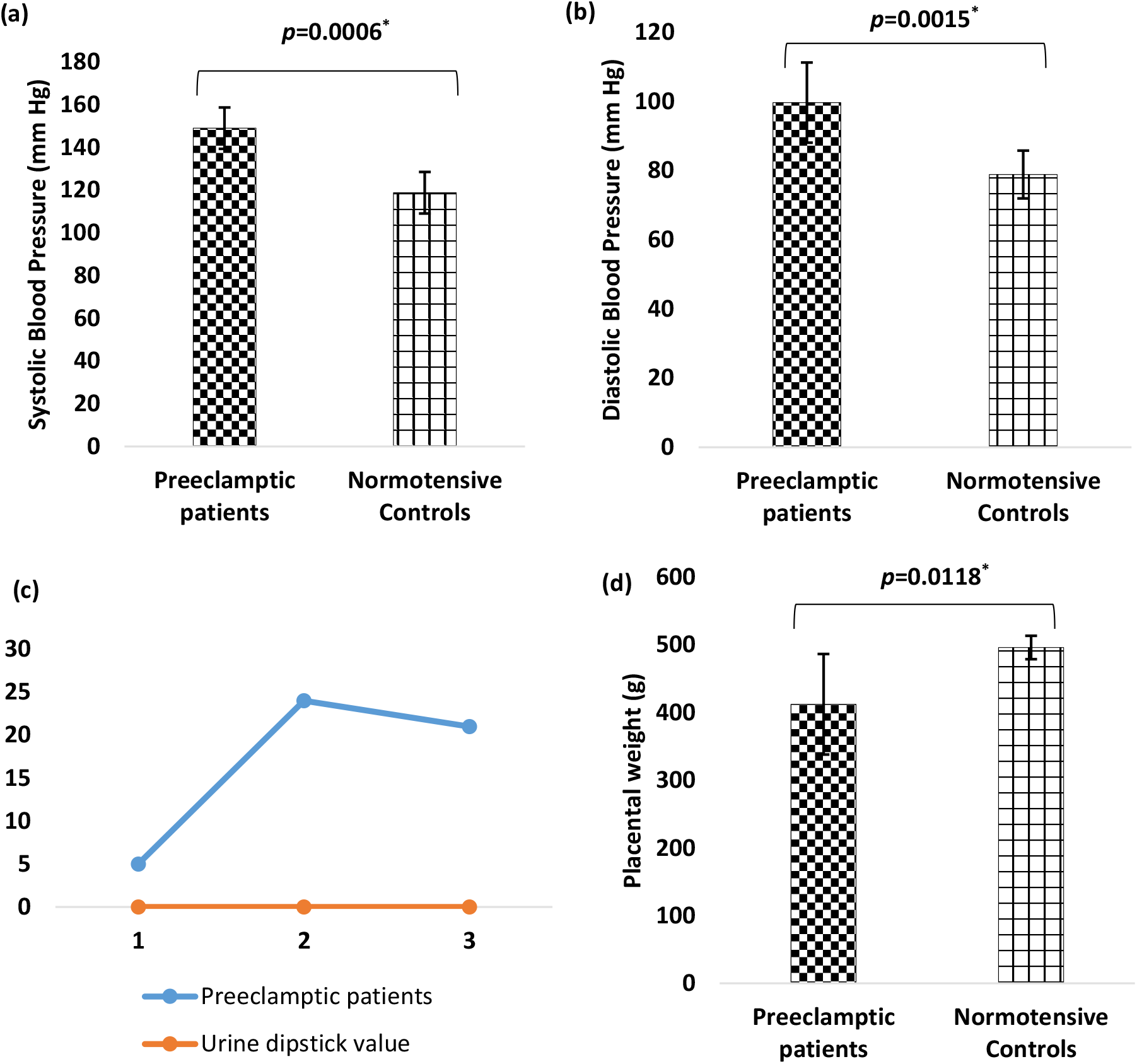
(a,b) Mean systolic and diastolic blood pressures in preeclamptic and control groups. (c) Line chart depicting trend of urine dipstick values in PE patients. (d) Placental weight in preeclamptic patients and controls. Paired t-test was used. Data expressed as Mean ± SD. *p*<0.05 was considered statistically significant.

### Pregnant women with preeclampsia manifested elevated levels of GRP78 and sFlt-1 protein

Circulating GRP78 levels in the maternal serum of preeclamptic patients (3044065±945822.8 pg/ml) were found upregulated as compared to controls (1983324±1180147 pg/ml) and the difference was statistically significant (*p*=0.0235). Circulating levels of sFlt-1 were also significantly raised (*p*=0.0131) in preeclamptic patients (29796.39±10070.01) as compared to normotensive, non-proteinuric pregnant women (controls) (17464.01±10373.32) (Figure 2 a,b).

**Figure 2:**
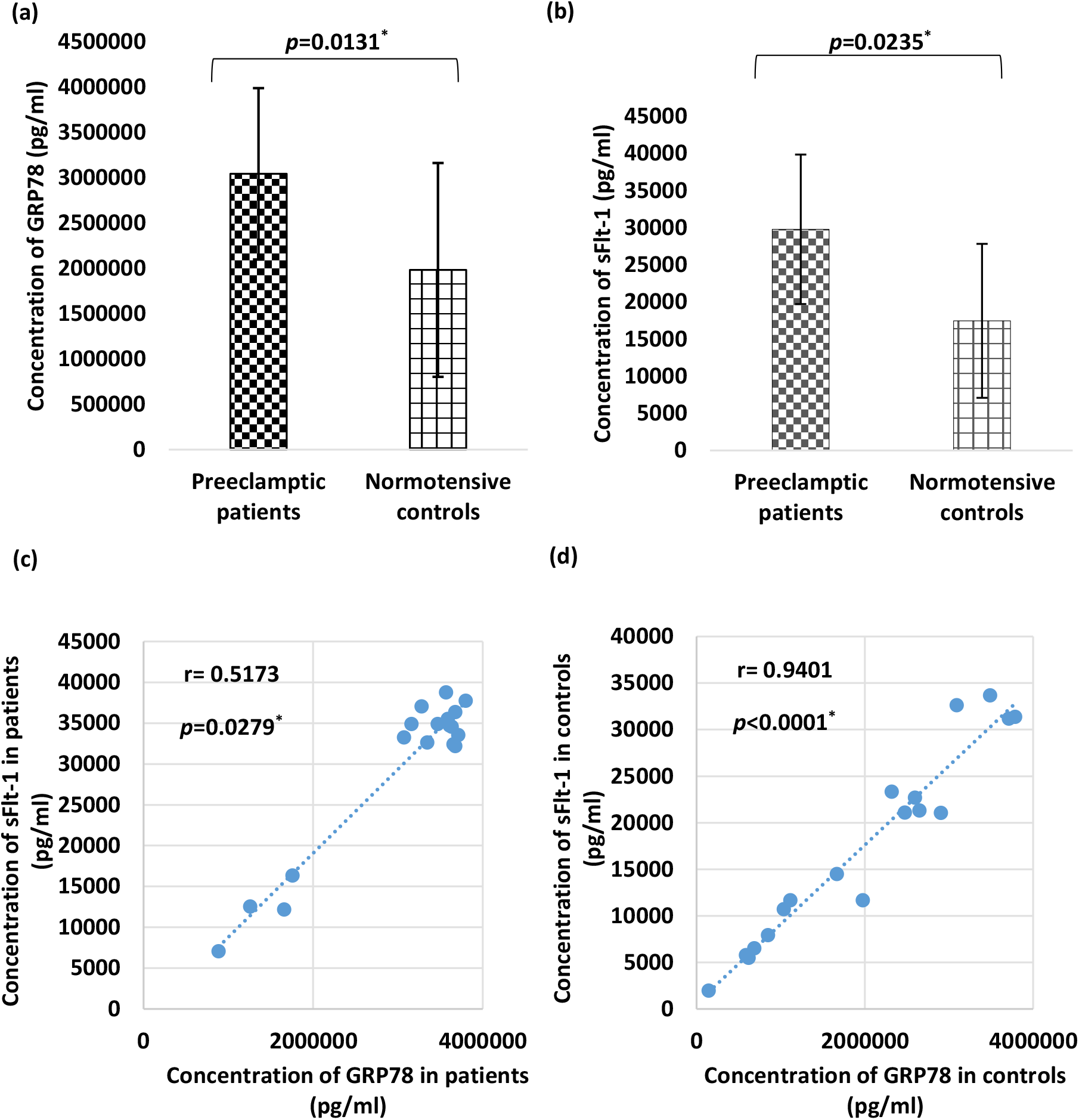
(a,b) Circulating levels of GRP78 and sFlt-1 in maternal serum of preeclamptic patients and controls (normotensive, non-proteinuric). Wilcoxon signed rank test was used for comparison. (c,d) Correlation analysis was done between GRP78 and sFlt-1 in patients and controls. Spearman correlation coefficient was used. Data expressed as Mean ± SD. *p*<0.05 was considered statistically significant.

### Correlation analysis

Significant positive correlation between circulating levels of GRP78 and sFlt-1 was observed in both patients and controls (Figure 2 c,d).

### Placentae of preeclamptic pregnancies demonstrated enhanced GRP78 protein expression

Immunofluorescence staining of placentae of preeclamptic mothers demonstrated stronger expression of GRP78 in syncytiotrophoblast and variable pattern from weak to moderate was seen around the walls of blood vessels and stromal cells in preeclamptic placentae (Figure 3b).

**Figure 3:**
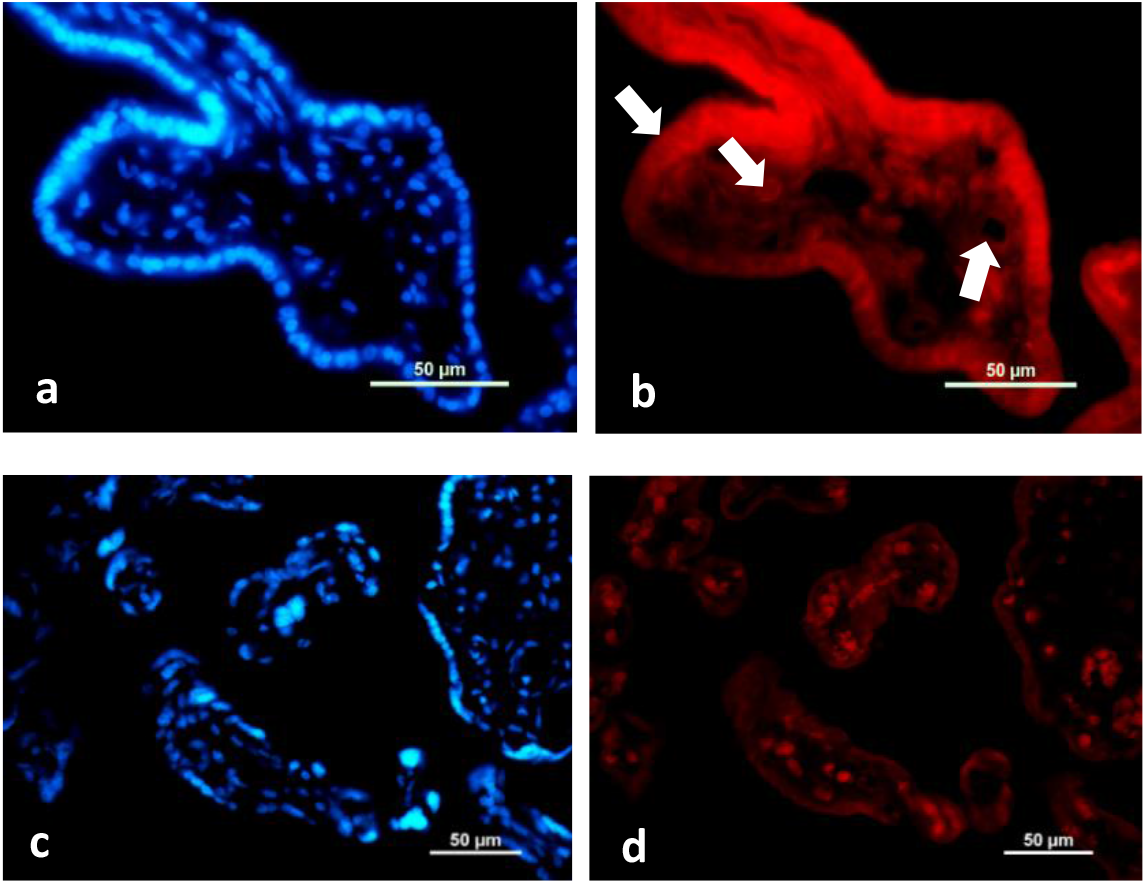
Representative immunofluorescence staining pattern of GRP78 protein in placentae from Preeclamptic (b) and Normotensive pregnant women (d). Positive staining (TRITC) for GRP78 is shown in syncytiotrophoblast and weak to moderate around wall of blood vessels and stromal cells. Arrows representing GRP78 positive cells (b). Nuclear staining (DAPI) is shown in blue (a,c).

### *In Vitro* experiments

### XBP1 profiling [Figure 4a-c]

### Immunofluorescence microscopy

Brighter signals of XBP1 protein (activated IRE1) at 14h were noted when BeWo cells were exposed to preeclamptic sera having raised concentrations of GRP78 and sFlt-1. Comparatively weaker signals were observed when cells were treated with normotensive sera. BeWo Cells when exposed to tunicamycin (5 μg/ml) demonstrated intense expression of XBP1 protein. Minimal expression of XBP1 protein was recorded in untreated BeWo cells. [Figure 4a].

**Figure 4:**
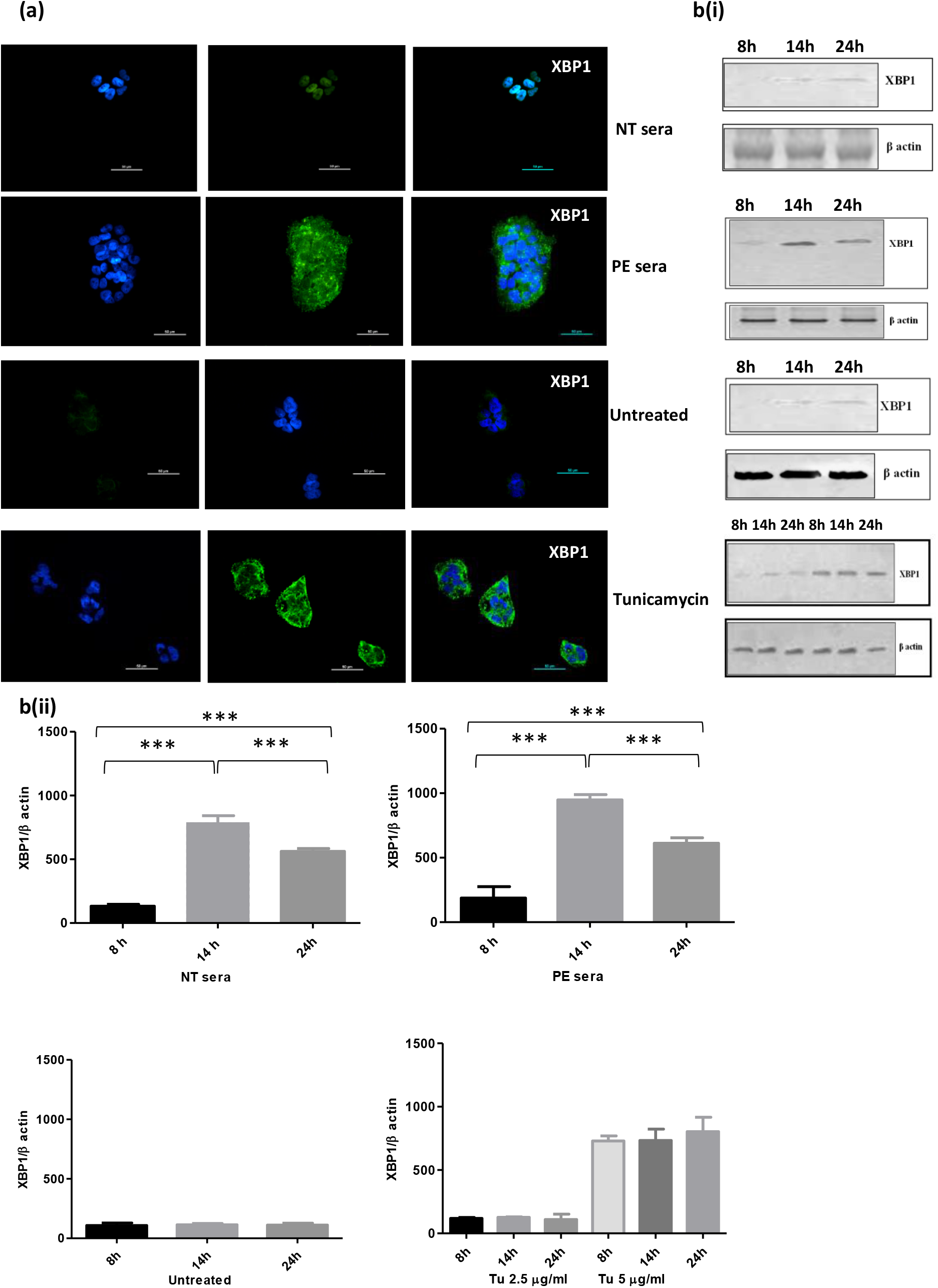
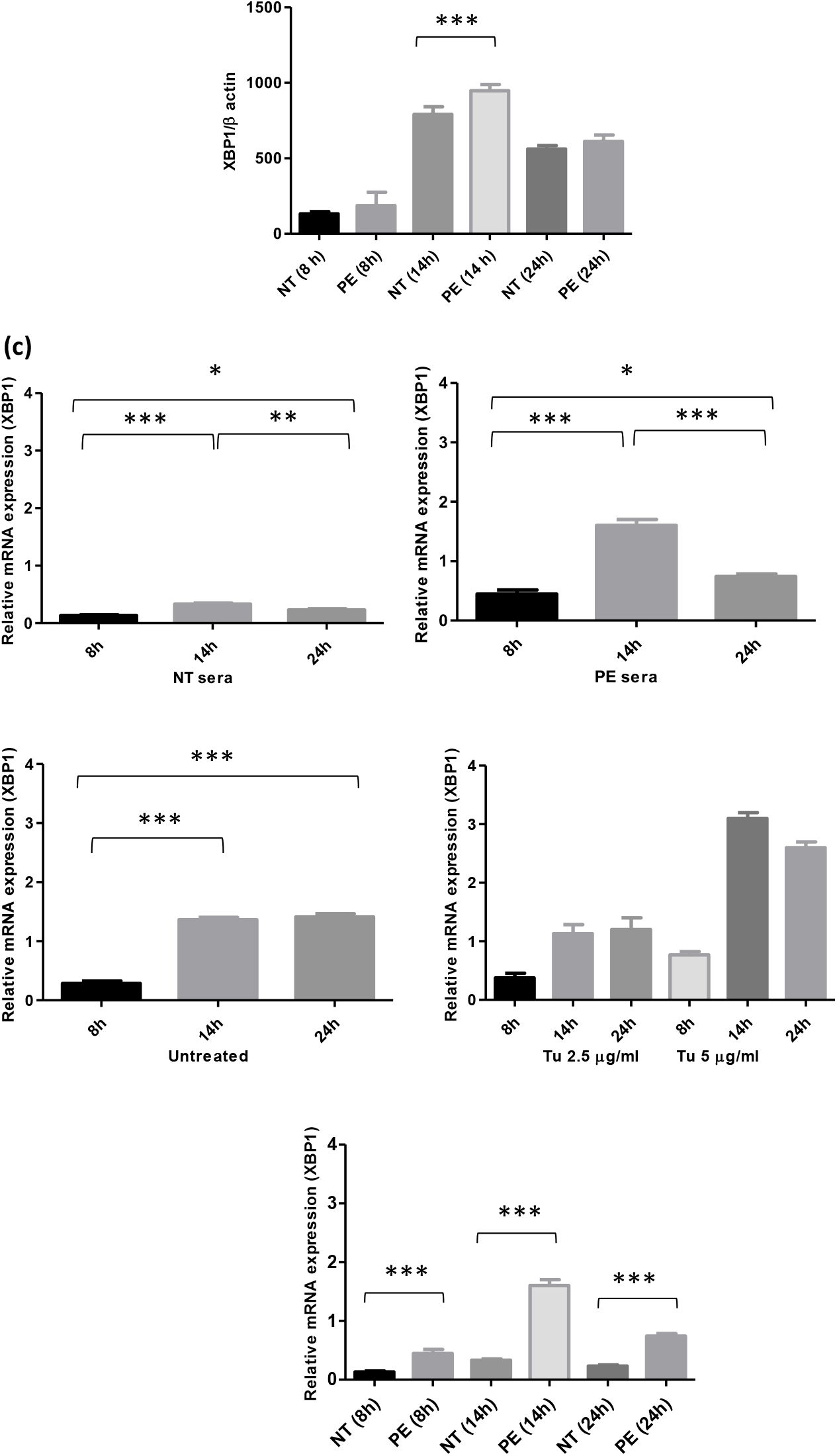
(a) Immunofluorescence staining pattern of anti-XBP1 antibody positive BeWo cells following 14 h treatment. b(i) Immunoblot representing expression of XBP1 in BeWo cells. β-Actin was used as protein loading control. b(ii) Bar diagrams represent normalized values. (c) Bar diagrams represent the relative mRNA expression of XBP1. β-Actin was used as positive control. Results are representative of 7 independent experiments. Data presented as mean ± SD. One way ANOVA with Bonferroni’s post hoc test was applied. *p* < 0.05 was considered statistically significant

### Western blot

Protein analysis of XBP1 marker by immunoblot manifested maximum XBP1 expression at 14h when BeWo cells were treated with preeclamptic sera (elevated levels of GRP78 and sFlt-1). Weaker expression was recorded when BeWo cells were exposed to normotensive sera with maximum signals at 24h. Tunicamycin (5 μg/ml) treated BeWo cells demonstrated enhanced protein expression of XBP1 at all-time points, as compared to its lower dose (2.5 μg/ml) [Figure 4b].

### mRNA expression (qRT-PCR)

mRNA expression was evaluated to validate the protein expression. XBP1 mRNA level was found maximum at 14h which was consistent with protein expression. Feeble mRNA expression of XBP1 was recorded in untreated BeWo cells [Figure 4c]. On comparing between the groups (1 and 2), enhanced mRNA levels of XBP1 (at all-time points, 8h 14h and 24h) was recorded for BeWo cells exposed to preeclamptic sera [Figure 4c].

### ATF6 profiling [Figure 5a-c]

### Immunofluorescence microscopy

Significant bright signals of ATF6 protein were observed at 14h in BeWo cells treated with preeclamptic sera. Weak expression of ATF6 was noted in cells treated with control sera (lower GRP78 and lower sFlt-1 concentration). As compared to control sera treated BeWo cells, cells treated with preeclamptic sera manifested enhanced signals of ATF6 protein. Intense expression of ATF6 was observed at 14 h in tunicamycin (5 μg/ml) treated BeWo cells. Very weak expression of ATF6 protein was demonstrated when BeWo cells left untreated [Figure 5a].

**Figure 5:**
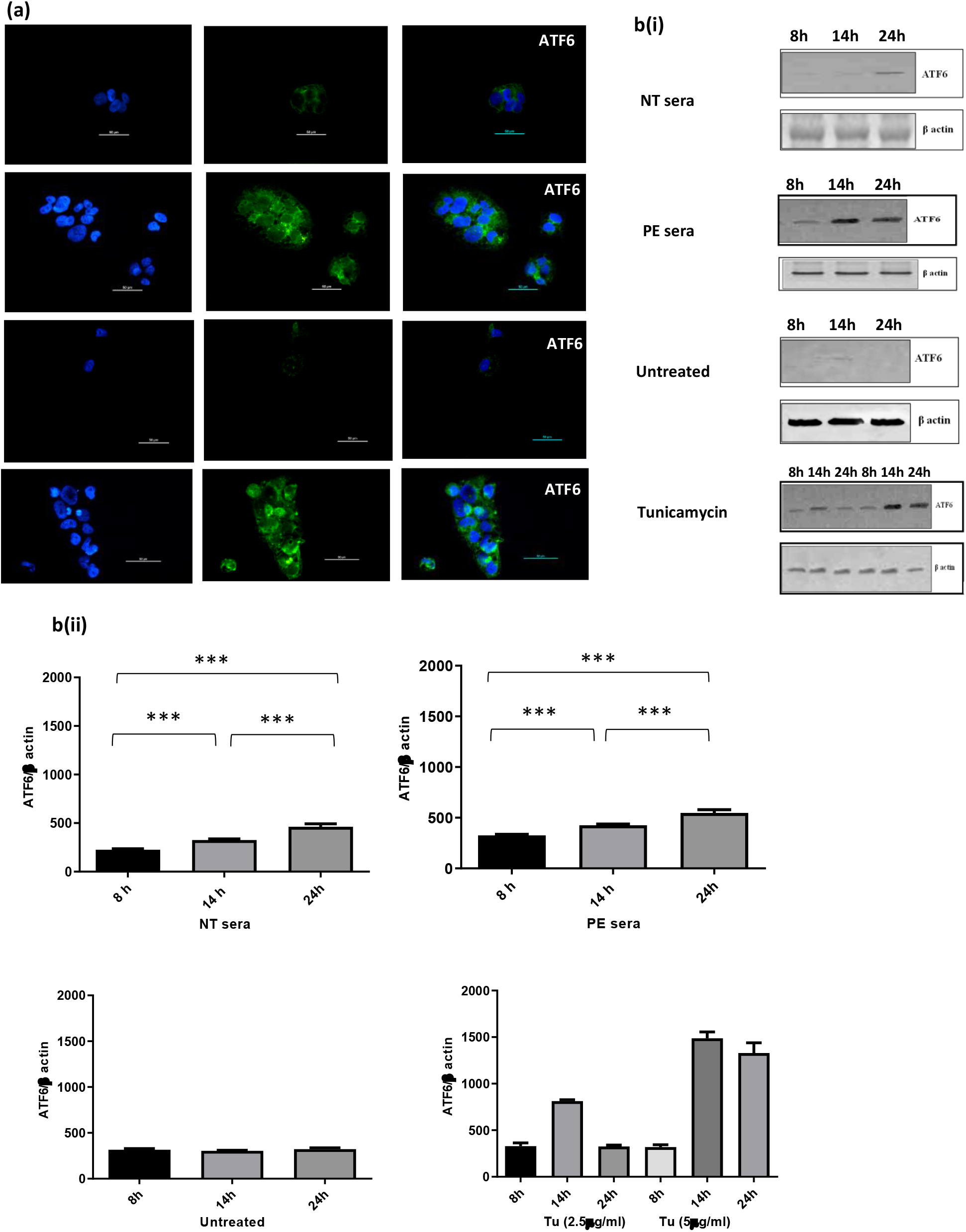
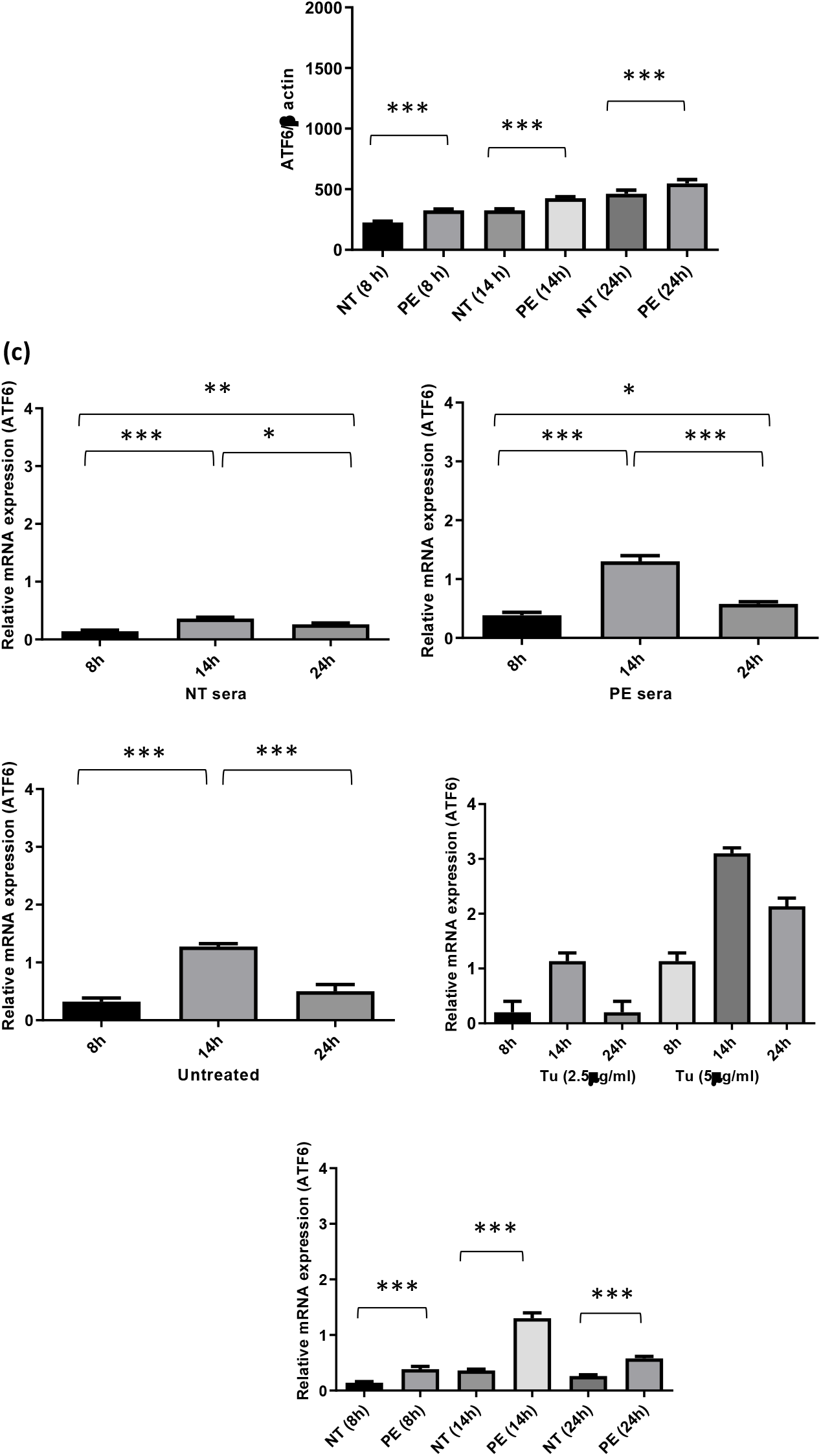
(a) Immunofluorescence staining pattern of anti-ATF6 antibody positive BeWo cells following 14 h treatment. b(i) Immunoblot representing expression of ATF6 in BeWo cells. β-Actin was used as protein loading control. b(ii) Bar diagrams represent the normalized values. (c) Bar diagrams represent relative mRNA expression of ATF6. GAPDH was used as positive control. Results are representative of 7 independent experiments. Data presented as mean ± SD. One way ANOVA with Bonferroni’s post hoc test was applied. *p* < 0.05 was considered statistically significant.

### Western blot

Maximum ATF6 protein expression was found at 24 h in both groups (1 and 2), however, expression was more in BeWo cells treated with preeclamptic sera, and the difference was statistically significant between the groups [Figure 5b].

### mRNA expression (qRT-PCR)

mRNA levels of ATF6 were found elevated at 14 h in both groups, 1 and 2. As compared to control sera treatment, ATF6 mRNA levels at all-time points, were enhanced in BeWo cells treated with preeclamptic sera and the difference noted was statistically significant [Figure 5c].

## Discussion

Apart from various stresses at cellular and subcellular level, placenta of preeclamptic pregnancies also undergo ER stress.^5,13^ The state of ER stress is triggered by conglomeration of mis-folded proteins in ER lumen as one of the canonical cellular stress pathways, collectively known as unfolded protein response (UPR).^8^ UPR, which primarily aims to restore the integrity of proteostatsis in ER lumen, comes into role when there is imbalance between unfolded/mis-folded proteins and folding capacity of the ER.^8^ Trafficking of translational proteins from cytosol to ER lumen is subjected to further folding, post-translational modifications and stabilization into their functional conformation.^14,15^ These functions are performed by ER folding machinery, composed of molecular chaperones, such as Glucose regulated protein78 (GRP78).^14,15^ GRP78, under normal state (in absence of stress) enchained with the luminal domains of three different ER-bound trans-membrane sensors; protein kinase-like endoplasmic reticulum kinase (PERK), inositol requiring enzyme (IRE1) and activating transcription factor (ATF6).^16^ The congregation of unfolded and/or mis-folded proteins calls upon GRP78 resulting in its dissociation from sensors and it is in this way three arms of UPR get activated leading to respective ER stress pathways.^16^

In the present study, we confirmed significantly raised levels of GRP78 in maternal serum as well as placentae of preeclamptic mothers as compared to normotensive controls (Figure 2a, 3b). Evidence from the literature states highly evolutionarily conserved pathways of UPR (IRE1 and ATF6) are either triggered secondary to perturbations in the ambience of endoplasmic reticulum (ie, ER stress) or independently, can modulate vascular growth.^17^ In accordance with aforesaid, negative modulation of vascular endothelial growth factor (with concomitant upregulation of sFlt-1) in placentae of preeclamptic mothers has already been reported in our previous studies.^18^ The up-regulation of sFlt-1 transcription as a consequence of placental ER stress may modulate maternal endothelial cell function and thereby contribute to the pathophysiology of PE.^19^ We, reported (amongst 50 pairs of patients and controls) significantly elevated levels of sFlt-1 in the preeclamptic patients as compared to controls. To decipher the pattern of levels of both GRP78 and sFlt-1, in maternal serum of preeclamptic mothers (n=50), we performed correlation analysis which revealed significant positive-correlation between GRP78 and sFlt-1, suggestive of PE patients (Spearman r = 0.5173; *p*= 0.0279*), who had higher GRP78 had also higher sFlt-1 in their sera when compared to controls (NT, NP). Till date, there is no evidence across the literature which could delineate whether GRP78 or sFlt-1 is the driving force and/or inducing agent to each other. We could not rule out the possibility of a presence of common /different factor(s) upregulating both GRP78 and sFlt-1 simultaneously/independently in preeclamptic subjects leading to activation of UPR and subsequently ER stress. Thus, there is scope of further experimentations to explore the intricacy of underlying mechanism.

The second part of study (*in vitro* experiments) was conducted to ascertain the combined effect of GRP78 and sFlt-1 present in maternal sera on IRE1 and ATF6 arms of UPR in trophoblast cells. BeWo cells (thus were chosen for further experimentations because these cells mimic *in vivo* syncytialisation of placental villous trophoblast) were exposed to preeclamptic sera having high GRP78 (2700000-2900000pg/ml) and high sFlt-1 (25000-27000pg/ml) levels. In the previous study, we have already reported the induction of PERK arm of UPR in BeWo cells.^20^ To confirm, whether or not, rest of the two pathways (IRE1 and ATF6) of UPR also get activated, BeWo cells were treated with elevated levels of GRP78 and sFlt-1, present in preeclamptic sera. To decipher the effect, signalling response of XBP1 (activated factor when IRE1 arm is induced) and ATF6 were observed at three different time points (8h, 14h, and 24 h) at protein and transcript levels.

The mammalian genome encodes two isoforms of IRE1, IRE1 α and IRE1β.^**21**^ The IRE1 α is expressed ubiquitously. A dissociation of IRE1α from GRP78 due to an elevated level of unfolded proteins initiate diverse downstream signaling of the UPR either through unconventional splicing of the transcription factor XBP-1 or/ and through post transcriptional modifications via Regulated IRE1-Dependent Decay (RIDD) of multiple substrates.^22-26^ The spliced XBP-1 enters into the nucleus to transcriptionally reprogram UPR target genes.^27^ In the present study, expression of XBP1 was up-regulated (maximum at 14 h) in BeWo cells treated with high GRP78 and sFlt-1 concentrations (PE sera) which reduced when treated with control sera. mRNA levels of XBP1 were also noted significantly upregulated in PE sera treated BeWo cells as compared to that of NT sera treatment, at all-time points and found consistent with that of protein expression. Our finding is consistent with the previous study by Ron *et al*. (2007) in terms of intensity of hourly expression of XBP1 after activation of IRE1 arm.^28^

Out of the two forms of mammalian homologous ATF6 proteins, ATF6α participates in the induction of various UPR target genes while ATF6β seems to have a minimal role in the UPR.^29,30^ ATF6 activation seems independent of IRE1, since its cleavage and the induction of GRP78 mRNA were unaffected in IRE1 null cells.^31^ Since long time ATF6 has been considered to fulfil merely adaptive functions during ER stress, and its sole intersection with ER stress-induced apoptosis consisted of a possible role in regulation of CHOP expression.^32^ In the present study, immunofluorescence microscopy revealed maximum ATF6 expression at 24h in BeWo cells exposed to control sera having low levels of GRP78 and sFlt-1, which is suggestive of adaptive efforts of ATF6 in maintaining proteostasis of ER machinery. Our result is in coherence with the finding of Thuerauf *et al*. (2004).^33^ As compared to control group, mRNA levels of ATF6 were significantly up regulated in PE sera at all-time points. So far, ATF6 arm is the most under investigated out of all the three arms of UPR. Its potential as an effector of clinical outcome is required to be explored further because ATF6, in particular, is one of the transcriptional regulators within the ER stress pathway.

## Summary

Preeclampsia has been recognized for at least 100 years however, its treatment has not changed significantly in over 50 years. Besides other agents contributing to preeclampsia, raised circulating levels of sFlt-1 has been a centre of attraction for nearly 2 decades. ER stress in preeclamptic placentae and in the trophoblast cells treated with preeclamptic sera may be the consequence of modulation of various post translational modifications of XBP1 and ATF6. Further pre-clinical, clinical investigations and validation will be required to determine the merits of UPR modulators as drugs against abnormal angiogenesis or to preserve endothelial function. This could go a long way in decreasing the maternal mortality in India if raised levels of sFlt-1 and expression of various ER stress markers are used adequately for screening of suspecting pregnant mothers for preeclampsia at the community level comprehensively.

## Authors’ contributions

SM and RD conceived and designed the study. SM did all the experiments. PA and SKG assisted in experiments. NB provided the clinical inputs. NR provided critical suggestions. SDD and SM did statistical analysis. SM and RD wrote first draft. RD and SM did compilation of the final draft.

