## Supplementary material for "Upregulated GRP78 and sFlt-1 in preeclampsia induces IRE1 and ATF6 arms of UPR leading to ER stress in placental cells": Tables

| Gene | Forward primer | Reverse primer |
|---|---|---|
| XBP1 | 5´ - TGGCCGGGTCTGCTGAGTCCG- 3´ | 5´ - ATCCATGGGGAGATGTTCTGG- 3´ |
| ATF6 | 5´ - CCACTAGTAGTATCAGCAGGAACTC- 3´ | 5´ - CCTTCTGCGGATGGCTTCAA- 3´ |
| GAPDH | 5´ - AGCCGAGCCACATC - 3´ | 5´ - TGAGGCTGTTGTCATACTTCTC - 3´ |
| β-Actin | 5´ - GAGCACAGAGCCTCGCCTTT - 3´ | 5´ - TCATCATCCATGGTGAGCTGG - 3´ |

Table1: Primers designed by NCBI

| **Study Groups** | | | |
|---|---|---|---|
| Clinical characteristics | Preeclamptic patients (n=50) | Controls (normotensive, non proteinuric) (n=50) | Statistical significance (*p* value)* |
| Systolic blood pressure (mmHg) | 148.99 ±9.68 | 118.77±9.73 | *p*=0.0006 |
| Diastolic blood pressure (mmHg) | 99.68±11.58 | 78.88±6.91 | *p*=0.0015 |
| Proteinuria (Urine dipstick test) | 5=1+, 24=2+, 21=3+ | Nil or traces | not applicable |
| Placental weight (g) | 412.22±74.26 | 496.11±17.30 | *p*=0.0087 |

Table2: Clinical Characteristics of Preeclamptic and Controls (normotensive, non proteinuric) pregnant women. n= number of subjects, Data presented as mean±SD, Paired t-test was used, *statistical significance, $p<0.05$.
